# Prototyping of a lateral flow assay based on monoclonal antibodies for detection of *Bothrops* venoms

**DOI:** 10.1101/2022.09.26.509137

**Authors:** Cecilie Knudsen, Jonas A. Jürgensen, Pelle D. Knudsen, Irina Oganesyan, Julian A. Harrison, Søren H. Dam, Aleksander M. Haack, Rasmus U. W. Friis, Selma B. Belfakir, Georgina M. S. Ross, Renato Zenobi, Andreas H. Laustsen

**Affiliations:** Department of Biotechnology and Biomedicine, Technical University of Denmark, Kongens Lyngby, Denmark; BioPorto Diagnostics A/S, Hellerup, Denmark; VenomAid Diagnostics ApS, Kongens Lyngby, Denmark; Department of Chemistry and Applied Biosciences, ETH Zürich, Zürich, Switzerland

**Keywords:** Snakebite envenoming, diagnostics, lateral flow assay, neglected tropical diseases

## Abstract

**Background:** Brazil is home to a multitude of venomous snakes, perhaps the most medically relevant of which belong to the *Bothrops* genus. *Bothrops spp*. are responsible for roughly 70% of all snakebites in Brazil, and envenomings caused by their bites can be treated with three types of antivenom: bothropic antivenom, bothro-lachetic antivenom, and bothro-crotalic antivenom. The choice in antivenom that is administered depends not only on its availability and how certain the treating physician is that the patient was bitten by a bothropic snake. The diagnosis of a bothropic envenoming can be made based on expert identification of a photo of the snake or based on a syndromic approach wherein the clinician examines the patient for characteristic manifestations of envenoming. This approach can be very effective but requires staff that has been trained in clinical snakebite management, which, unfortunately, far from all relevant staff has.

**Results:** In this paper, we describe a prototype of the first lateral flow assay (LFA) capable of detecting venoms from Brazilian *Bothrops spp*. The monoclonal antibodies for the assay were generated using hybridoma technology and screened in sandwich enzyme-linked immunosorbent assays (ELISAs) to identify *Bothrops spp*. specific antibody sandwich pairs. The sandwich pairs were used to develop a prototype LFA that was able to detect venom from several different *Bothrops spp*. The limit of detection (LoD) of the prototype was evaluated using Brazilian *B. atrox* whole venom and was determined to be 8.0 ng/mL in spiked serum samples and 9.5 ng/mL in spiked urine samples, when using a portable reader, and < 25 ng/mL in spiked buffer when reading by eye.

**Significance:** The work presented here serves as a proof of concept of a genus-specific venom detection kit, which could support physicians in diagnosing *Bothrops* envenomings. Although further optimization and testing is needed before the LFA can find clinical use, such a device could aid in decentralizing antivenoms in the Brazilian Amazon and help ensure optimal snakebite management for even more victims of this highly neglected disease.

## 1 Introduction

Snakebite envenoming has plagued mankind since time immemorial, annually exacting a toll of 81,000-138,000 deaths and roughly six million disability-adjusted life-years (DALYs) [1,2]. Brazil alone experiences approximately 26,000-30,000 snakebite envenomings per year, most of which are caused by *Bothrops* species, with the species *B. atrox* contributing with an especially high number of bites [3–5]. In addition to the *Bothrops* genus, bites from *Crotalus durissus, Lachesis muta*, and *Micrurus* spp. also occur, although to lesser extents [3]. The mainstay treatment of snakebite envenoming is antivenom, and, in Brazil, antivenoms are available at a genus and inter-genus level in the form of bothropic antivenom, crotalic antivenom, bothro-lachetic antivenom, bothro-crotalic antivenom, and elapidic antivenom. The corollary is that snakebite diagnosis in Brazil must be undertaken at the same level; healthcare providers must determine whether the bite was dry (i.e., a bite in which no venom was injected), whether it warrants antivenom administration, and, if so, which antivenom is appropriate when factoring in the snake that caused the bite [6,7]. This is usually accomplished via a syndromic approach, where the clinical manifestations of envenoming are compared to those associated with the different snake genera [6]. Unfortunately, some of the syndromes overlap, as the venoms of most Central and South American pit vipers are known to cause similar local effects and coagulopathies [6,8]. The venom of South American *Crotalus* species might be the exception, as they typically cause only mild local effects and have instead been shown to cause more severe systemic effects, including neurotoxicity [6]. The syndromes of envenomings caused by *Bothrops* and *Lachesis* species are even more similar, thus further complicating correct diagnosis, but can in some cases be distinguished by the vagomimetic effects of lachetic venom on the gastrointestinal system. However, the presence of such effects cannot be used to confirm a lachetic envenoming, equally, their absence cannot be used to exclude it [6]. These overlapping syndromes can complicate matters for healthcare providers trying to identify the optimal treatment, especially as not all healthcare providers have received adequate training in clinical snakebite management [9]. This is unfortunate, as early and correct treatment has been shown to correlate with better patient outcomes for snakebite victims both in Brazil and abroad [10–14].

To facilitate early and correct treatment of snakebite envenoming in Brazil, the development of supportive diagnostic tools capable of distinguishing Brazilian pit viper bites at the same level at which clinical decisions are made (e.g., the genus level) could be beneficial. Such diagnostic tools could support efforts to secure faster treatment in the Amazon region, e.g., through antivenom decentralisation. Empowering healthcare providers at remote clinics to diagnose snakebites more easily and with higher precision enables them to choose the appropriate antivenom to treat envenomings [15]. In more metropolitan areas of Brazil, snakebite victims do not have to travel as far to reach a healthcare facility, and thus might present to the hospital before clinical manifestations of envenoming develop. In such scenarios, diagnostic tools might be able to speed up the diagnosis by providing healthcare workers with an idea of the offending snake before the venom has exerted its full toxic effects in the patient, and, as such, would allow healthcare workers to prepare the correct treatment regime early on. Finally, snakebite diagnostic tools might also help lower the incidence of misdiagnosis of snakebite patients. Here, we describe an attempt to develop such a snakebite diagnostic tool in the form of a lateral flow assay (LFA). LFAs are well-suited for the diagnosis of snakebite envenoming for several reasons. E.g., the widespread use of LFAs during the COVID-19 pandemic has demonstrated the feasibility of implementing such rapid diagnostics in both larger clinics, more remote settings, and even for home use by consumers. Moreover, LFAs are affordable, rapid, and user-friendly requiring no specialised equipment or training to operate them. Combined, this makes LFAs suitable for point of care (PoC) use, especially in primary care settings. Additionally, they can be mass produced for as little as 0.10-3.00 USD per test [16]. The affordability of these tests is especially relevant, as snakebite envenoming is associated with poverty and often occurs in regions with low-resource healthcare systems, meaning that the cost of treatment and diagnostics could easily become prohibitively high [17–19]. Finally, LFAs can be carried out in 5-20 minutes, making them appropriate diagnostics for an acute disease, such as snakebite envenoming. An additional advantage, is that LFAs rely on paper-based, disposable materials, making them relatively sustainable compared to diagnostics requiring more plastic components and/or harmful chemicals (e.g., enzyme-linked immunosorbent assays, abbreviated ELISAs). In this work, specifically, we have developed a prototype sandwich LFA using monoclonal antibodies capable of detecting venom from multiple Brazilian *Bothrops* species without cross-reacting with venoms from other Brazilian vipers. If taken into clinical development, such a tool could support clinical diagnosis and decision-making regarding treatment, thereby contributing to more snakebite patients in Brazil receiving the antivenom most appropriate for their envenoming as early as possible.

## 2 Materials & Methods

### 2.1 Immunisation

Lyophilised whole venom from *Bothrops atrox* (a specimen from Brazil, Latoxan, L1210A) was reconstituted in a sterile 0.9% saline solution to a final concentration of 1 mg/mL. Either 5 or 10 µg venom, depending on the protocol, were mixed with aluminium hydroxide at a ratio of 1 mg aluminium hydroxide per 25 µg venom, in a solution of 0.05% methiolate, 50% Adjuvant P (Gerbu, 3111.6001), and 0.9% saline water to a volume of 100 µL to create an immunisation mixture. Immunisation mixtures were injected subcutaneously into four Naval Medicinal Research Institute (NMRI) mice: Two mice were injected with 5 µg venom, and two were injected with 10 µg venom. The mice were injected on days 1, 14, and 28, and they were bled on days 25 and 38 by taking a sample of 100 µL blood from the jaw of each mouse and transferring it to tubes containing EDTA. Subsequently, 200 µL 0.9% saline solution was added to the blood sample and the solution was centrifuged at 2,000 g for five minutes at room temperature, enabling the extraction of 250 µL plasma, which was tested in ELISA as described below to monitor antibody development. On day 42, the animals received an immunisation boost consisting of 20 µg whole venom dissolved in 0.9% saline to a volume of 100 µL, injected intravenously.

### 2.2 Bleed screenings

Clear 96-well plates (Thermo Scientific, MaxiSorp, 439454) were coated with 100 µL/well of 1000 ng/mL whole venom from either *B. atrox, L. m. muta*, or *C. d. terrificus* dissolved in carbonate buffer (made from tablets as per the manufacturer’s instructions, Medicago, 09-8922). The plates were incubated overnight at 4 °C or at ambient temperature and shaken at 300 rpm for two hours, before being washed thrice with washing buffer (Ampliqon Laboratory Reagents, AMPQ40825.5). Afterwards, 110 μL of mouse plasma diluted 1:55 in dilution buffer (10 mM phosphate, 140 mM NaCl, 0.5% w/v BSA, 0.0016% w/v phenol red, 0.05% v/v Tween-20, 0.1% v/v ProClin 950, pH 7.4) were added to each well. The plates were left shaking at 300 rpm for 1 hour, and then washed thrice with washing buffer. Next, 100 μL of a 1:1000 dilution of horseradish peroxidase (HRP)-conjugated, polyclonal, rabbit anti-mouse antibody (Dako, P0260) in dilution buffer (final concentration 1.3 ng/mL) was added to each well. The plates were left shaking at 300 rpm for 1 hour and subsequently washed thrice with washing buffer. Then, 100 μL of 3,3’, 5,5’ tetramethylbenzidine (TMB) One substrate (Eco-Tek, 4380-12-15) were added to each well, and the plates were left in complete darkness for 12 minutes, before 100 μL of 0.5 M sulphuric acid were added to all wells to stop the reaction. The absorbance was measured at 450 nm and 620 nm on a Thermo Multiscan Ex plate reader.

### 2.3 Cloning and hybridoma generation

On day 45 of the immunisation schedule, the mice were sacrificed, and their spleens were extracted. The spleens were reduced to a single-cell-suspension with a mortar and pestle and were immediately mixed with SP2/0-AG14 myeloma cells and a PEG solution to fuse the B cells from the spleen with the myeloma cells. The resulting cells were spread into 96-well microtiter plates and grown in HAT medium for 7-10 days to select successfully fused cells. Culture supernatant from the different wells were tested with ELISA as described above, with the exceptions that an IgG-specific HRP-conjugated antibody (Merck, AP127P) was used for detection, and instead of mouse plasma, 1:50 and 1:100 dilutions of supernatant from the hybridoma growth media were used. The cell cultures corresponding to wells with positive ELISA signals for *B. atrox* venom and negligible signals for *C. d. terrificus* and *L. m. muta* venom were selected for cloning (other signal combinations were also chosen for completeness). These cells were transferred to HT medium and cultured further. The cells were then sequentially diluted in 96-well microtiter plates and tested with ELISA as described above, until wells with single cells were identified. These monoclonal cell lines were expanded and used in future experiments.

### 2.4 Antibody purification

NaCl was added to antibody-containing hybridoma culture supernatant to a final concentration of 2.5 M. After the salt was dissolved, half a teaspoon of Celpure P65 (Honeywell, 525235) was added, and the culture supernatant was filtered through a 0.45 µm filter (Durapore® Membrane Filters 0.45 µm, HVLP04700). An ÄktaPrimePlus system was washed with ultrapure water to remove air in the tubing. The system was then primed with filtered culture supernatant, and the purification procedure was started (rProtein A sepharose Fast Flow (GE healthcare: 17-1279-03)), and the antibodies were purified.

### 2.5 Cross-reactivity screenings

The antibody binding profiles were investigated by screening the antibodies in ELISA against whole venom from *B. atrox, L. m. muta*, and *C. d. terrificus*. Antibodies that recognised only *B. atrox* venom were screened in ELISA against a panel of 21 venoms from the following species: *B. alternatus, B. asper, B. atrox* (Brazil), *B. atrox* (Columbia), *B. atrox* (Suriname), *B. jararaca, B. jararacussu, B. leucurus, B. mattogrossensis, B. moojeni, B. neuwiedi diporus, B. neuwiedi neuwiedi, B. pauloensis, C. adamanteus, C. atrox, C. d. terrificus, C. horridus, C. scutulatus scutulatus, C. simus, L. melanocephala*, and *L. m. muta* the venoms used in this study are summarised in the **Supplementary Information (SI) Table S1**. The ELISAs were carried out as described for the bleed screenings, with the exception that 100 µL/well purified antibodies in dilution buffer at 1000 ng/mL were used instead of mouse serum. Later, these cross-reactivity screenings were repeated with sandwich ELISAs and LFAs, using the protocols described below. Additionally, the following venoms were screened with LFA: *Agkistrodon bilineatus howard gloydi, Atropoides mexicanus, A. picadoi, B. andianus, B. barnetti, B. castelneudi, B. chloromelas, B. hyoprorus, Bothriechis lateralis, B. microphtalmus, B. peruviensis, B. pictus, B. schlegeli, Cerrophidion sasai, L. stenophrys, Porthidium nasutum*, and *P. ophryomegas*.

### 2.6 Biotinylation of antibodies

The antibodies were buffer exchanged into carbonate buffer using Nap5 columns (Illustra, GE Healthcare, 17085302) according to the manufacturer’s protocol. The antibodies were eluted from the Nap5 columns with 1.5 mL of carbonate buffer, and the eluate was collected into a Vivaspin 6 50 kDa MWCO column (Sartorius, V10631), which was centrifugated at 2,113 g for five minutes to concentrate the antibodies. The antibody concentration was determined using an Eppendorf BioPhotometer model 6131. Biotin-N-hydroxysuccinimide (Sigma-Aldrich, H1759-5mg) was dissolved in DMSO to a concentration of 0.4 µg/mL. Biotin-N-hydroxysuccinimide was added to the buffer exchanged antibodies at a ratio of 55 µg Biotin-N-hydroxysuccinimide per mg antibody, and the samples were vortexed immediately for one minute, before being left with end-over-end rotation for two hours. The reaction was stopped through addition of 50 µL 1 M Tris per 2.5 mL sample. The antibodies were buffer exchanged into 0.14 M PBS with 0.1% NaN_3_, concentrated, and their concentrations were measured again. The biotinylated antibodies were evaluated in ELISA as described previously, with the exception that antibody dilution series in the range 0.5-500 ng/mL were made of the biotinylated and unbiotinylated antibodies, respectively, using dilution buffer. The biotinylated and unbiotinylated antibodies in the dilution series were detected with different reagents: The biotinylated antibodies were detected with HRP-conjugated streptavidin (for two replicate dilution series) and with a 1:1000 dilution of HRP-conjugated anti-mouse antibody (Dako, P0260) (for another two replicate dilution series), while the unbiotinylated antibodies were only detected with the 1:1000 dilution of the HRP-conjugated anti-mouse antibody (also for two replicate dilution series). The HRP-conjugated streptavidin used for detection was prepared by reconstituting lyophilised HRP-conjugated streptavidin (KemEnTec, 14-30-00) in 50% glycerol and leaving it with end-over-end rotation for at least 90 minutes, before being diluted 1:5000 in HRP-StabilPlus buffer (KemEnTec, 4530A).

### 2.7 Sandwich pair screening

100 µL of 1000 ng/mL of unbiotinylated antibodies in carbonate buffer were coated in individual wells on 96-well plates (Thermo Scientific, MaxiSorp, 439454) and incubated at 4 °C overnight or for two hours at ambient temperature and 300 rpm. The plates were washed thrice with washing buffer, 100 µL of 1000 ng/mL whole venom dissolved in dilution buffer were added to each well, and the plates were incubated for another hour at ambient temperature and 300 rpm. The plates were washed thrice, and 100 µL of 1000 ng/mL biotinylated antibody in dilution buffer were added to each well, before the plates were incubated and washed again thrice. From here on, the protocol is identical to those previously described with HRP-conjugated streptavidin used as a detection reagent.

### 2.8 Gold-conjugation of antibodies

The antibodies were buffer exchanged into ultra-pure water, concentrated, and the concentration was measured as described above for biotinylations. The antibodies were gold-conjugated using 40 nm gold particles at 15 OD/mL from a Naked Gold Conjugation Kit (BioPorto Diagnostics, NGIB18) according to the manufacturer’s protocols. In addition to the salt tests described in the manufacturer’s protocol, the suitability of the gold-conjugated antibodies was also evaluated in terms of false positives by comparing the results of positive and negative samples on LFA strips (see below).

### 2.9 Lateral Flow Assays

75 µL running buffer (0.14 M PBS with 50 g/L BSA, 0.5% Tween-20, 0.1% ProClin 950) were added to a tube and mixed with 5 µL 0.1 mg/mL biotinylated capture antibody (dissolved in running buffer), 5 µL gold-conjugated detection antibody, and 15 µL sample. The mixture was allowed to incubate for five minutes at room temperature. After 5 minutes, a commercially available LFA strip (BioPorto Diagnostics, gRAD1-120) was inserted into the tube. The strip was then either read every 10 seconds for 15 minutes (kinetic measurements) or read once after 15 minutes (point measurement) using a Cube Reader (ChemBio Diagnostics). Samples consisted of either running buffer, pooled human serum (Sigma-Aldrich, H4522-100mL), or pooled human urine (Lee Biosolutions, 991-03-P) spiked with whole venom. For the interference study, each interferant was diluted in the matrix (pooled human serum or pooled human urine) to the final concentration seen in **Error! Reference source not found**.. The measurements were analysed in GraphPad Prism 9 (version 9.4.0) using a one-way ANOVA analysis followed by a post-hoc Dunnett analysis comparing each interferant mean to the mean of the control.

### 2.10 Lyophilisation

Lyoprotectant solutions were prepared, which consisted of 0.3% w/v of either BSA or casein and 5% w/v of either trehalose, sucrose, or mannitol, dissolved in ultrapure water. Gold-conjugated antibodies were dissolved in these solutions to a final concentration of 10% v/v along with biotinylated antibodies with a final concentration of 10 µg/mL. The mixtures were aliquoted 50 µL at a time into 0.5 mL tubes and lyophilized at -40 °C in a lyophiliser (Labogene Scanvac superior touch 55-80) for approximately 12 hours. After lyophilisation, the tubes were sealed and stored at 4 °C until use. For the stability studies, the samples were used the day after the lyophilisation.

### 2.11 Sample preparation for native mass spectrometry

*B. atrox* venom and antibody samples were fractionated and exchanged into 200 mM ammonium acetate by size exclusion chromatography (SEC) as previously described [20,21]. These experiments were performed on a Superdex Increase 200 10/300 GL column (Cytiva, Massachusetts, United States) pre-equilibrated with 200 mM ammonium acetate at the rate of 0.5 mL/min. Samples were collected and stored a 4 °C until used. The concentration of toxins in the SEC fractions was not adjusted prior to mixing with the antibody.

### 2.12 Native mass spectrometry

All mass spectrometry (MS) experiments were performed on a SELECT SERIES cyclic IMS mass spectrometer (Waters, Manchester, U.K.), which was fitted with a 32,000 m/z quadrupole, equipped with an electron capture dissociation (ECD) cell (MSvision, Almere, Netherlands) in the transfer region of this mass spectrometer. Approximately 4 µL of sample were nano-sprayed from borosilicate capillaries (prepared in-house) fitted with a platinum wire. Spectra were acquired in positive ion mode, with the *m/z* range set to 50-8,000. Acquisitions were performed for five minutes at a rate of 1 scan per second. The operating parameters for the MS experiments were as follows: capillary voltage, 1.2 - 1.5 kV; sampling cone, 20 V; source offset, 30 V; source temperature, 28 °C; trap collision energy, 5 V; transfer collision energy, 5 V; and Ion guide RF, 700 V. This instrument was calibrated with a 50:50 acetonitrile:water solution containing 150 µM caesium iodide (99.999%, analytical standard for HR-MS, Fluka, Buchs, Switzerland) each day prior to measurements.

## 3 Results

### 3.1 Antibody discovery and characterisation

Mice were immunised with whole venom from Brazilian *B. atrox* specimens. ELISAs on the plasma from the mice were used to confirm that venom-specific antibodies had been raised. Once the immunisation schedule had been completed, the mice were euthanised, and B cells were harvested from their spleens. Hybridoma cell lines were generated, screened for expression of antibodies specific to venom, and cloned to monoclonality. This resulted in 38 monoclonal cell lines expressing antibodies capable of recognising *B. atrox* venom. The 38 resulting antibodies were screened in ELISAs against whole venom from *B. atrox, C. d. terrificus*, and *L. m. muta*, to identify antibodies that bind only to *Bothrops* venom, without cross-reacting to venoms from either of these two other medically relevant pit vipers. Out of these 38 antibodies, four bound only *B. atrox* venom (*B. atrox* signal > 3.0, other signals < 0.5). These four antibodies were screened in further ELISAs against a panel of 21 Latin American snake venoms (including the original three venoms), and three out of the four antibodies were selected, as they retained a binding profile where only *Bothrops* venoms elicited strong signals (*Bothrops* signal > 2.5, other signals < 0.5). The fourth antibody, conversely, was shown to also react with *C. horridus* venom (**Error! Reference source not found**.).

The antibodies that had been selected based on their binding profiles were biotinylated and screened against all 38 monoclonal antibodies in sandwich ELISAs to find the sandwich pairs necessary for the LFA. Our hypothesis was that only one sandwich pair component would need to have the desired genus specificity for the pair to be genus-specific. With the three selected detection antibodies, we found ten possible sandwich pairs that could detect *B. atrox* venom. All these pairs utilised the same detection antibody (antibody 86-14), while no sandwich partners were found for the other two detection antibodies. We subsequently investigated the binding profiles of these ten sandwich pairs against the panel of 21 venoms and confirmed that the binding profiles of the sandwich pairs reflected the binding profile of the genus-specific detection antibody (**Error! Reference source not found**.). Out of the ten sandwich pairs, four seemed especially interesting due to the combination of their recognition of venoms from multiple *Bothrops* species and the comparatively high signals they yielded in the sandwich ELISAs (**Error! Reference source not found**.). These four pairs were selected for further evaluations in LFAs.

### 3.2 LFA prototyping

The four selected non-genus-specific capture antibodies were biotinylated as preparation for use in LFAs, and the genus-specific detection monoclonal antibody was conjugated to gold nanoparticles (AuNP-mAb). The gold-conjugation was carried out at ten different pHs, and the absorption spectra of each version of the conjugated antibody were investigated. Additionally, the blank signal (i.e., the test line signals of LFAs in which no antigen was added) of the different AuNP-mAbs were measured in LFAs to find the optimal conjugation conditions. Following successful conjugation, LFA prototypes were established for each of the four sandwich pairs using Generic Rapid Assay Device (gRAD) LFA strips [22]. The gRAD is a commercially available, universal sandwich LFA that is not antigen-specific. The gRAD’s single test line is composed of biotin-binding proteins and its control line is immobilised anti-mouse antibodies. This generic configuration makes it possible to rapidly prototype LFAs, as long as the capture antibody is biotinylated and the detection antibody is murine. The gRAD-based prototypes were used to detect a 1000 ng/mL solution of whole venom from *B. atrox* (this concentration is higher than the expected levels detected after a bite [23,24], and was intended as a positive control), and the test line and control line intensities were quantified using a low-cost commercial LFA reader. Out of the four prototypes, the one relying on antibodies 86-14 and 86-11 provided the highest test line signal, so it was decided to keep working with this prototype. The prototype’s signal-dependence on the antigen concentration was assessed by measuring a dilution curve consisting of LFA running buffer spiked with various concentrations of *B. atrox* whole venom. While there was a clear correlation between venom concentration and test line signal intensity, the correlation was not linear (see **Figure 1A, Figure 1B** and **Error! Reference source not found**.). The prototype was also used to measure LFA running buffer spiked with higher concentrations (5,000 - 5,000,000 ng/mL) of *B. atrox* whole venom to assess whether the prototype is affected by the hook effect (also known as prozone effect) (see **Figure 1C** and **Figure 1D**). The prototype appears to be influenced by high antigen concentrations, as at concentrations above 20,000 ng/mL, the test line signal starts decreasing, even as the antigen concentration increases. This reduction of test line intensity at high antigen concentrations is characteristic of the hook effect [25]. Finally, the visual (by naked eye) and digital (by low-cost reader) limit of detection (LoD) of the prototype was determined and it was assessed how this LoD might be influenced by the sample matrix. Therefore, LFA running buffer, pooled human serum, and pooled human urine were spiked with 0-20 ng/mL of *B. atrox* whole venom, and the test and control line signal intensities were measured. Linear regression was used to find the linear functions that best described the data. The LoDs were calculated using the formula LoD = 3.3*(σ/S), where σ is the standard deviation of the test line signal, S is the slope of the curve, and 3.3 is a constant [26]. The parameters used in the calculations and the resulting LoDs can be seen in **Table 1**, while the test line signals are plotted in **Figure 1E** and **Figure 1F**. Spiking LFA running buffer and urine with venom generally resulted in lower false positive signal (i.e., test line signal on tests in which no antigen was added) than did spiking serum with venom. However, the curve based on the serum samples had a steeper slope than the curves based on the LFA buffer and urine samples. The LoD was therefore determined to be lower in serum samples (8.0 ng/mL) than in urine samples (9.5 ng/mL) and LFA buffer samples (10.3 ng/mL). The visual LoD (i.e., the lowest antigen concentration that could be read by the naked eye) was slightly higher, i.e., a visual LoD of 25 ng/mL was comfortably achievable (**Error! Reference source not found**.), indicating that the test is usable even without any specialised LFA reader.

**Table 1.**
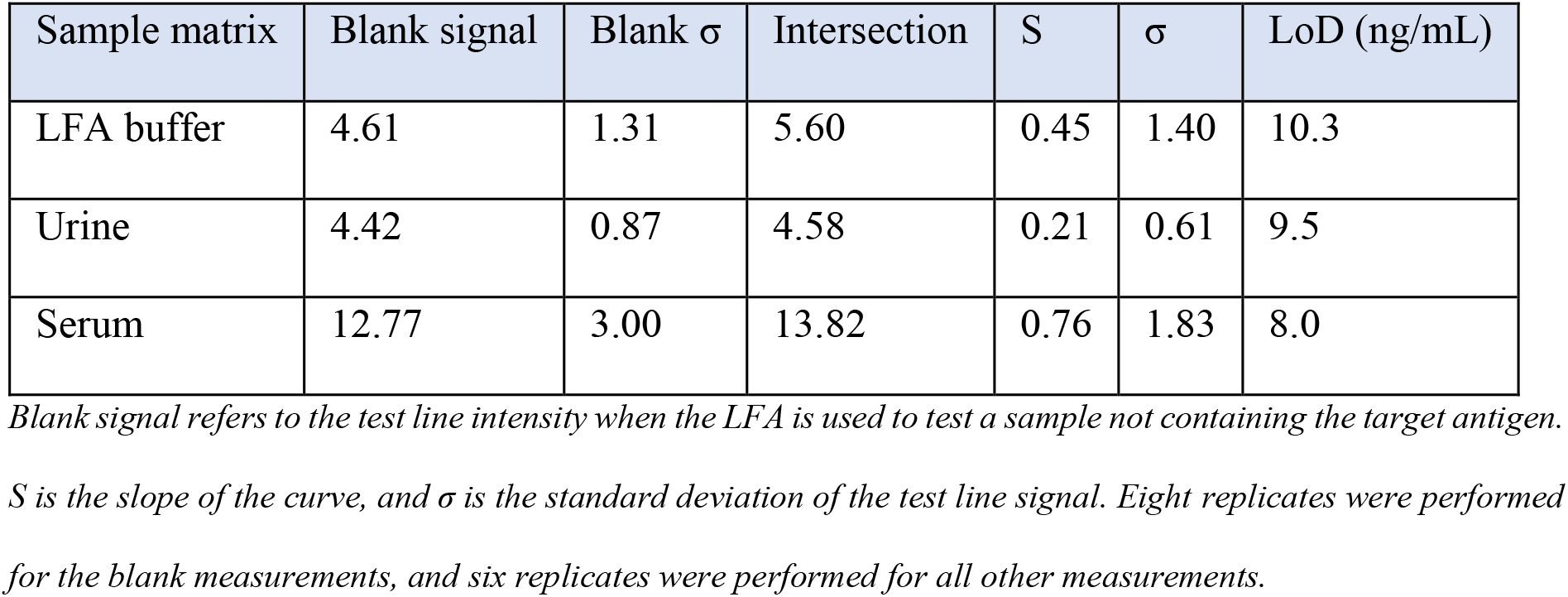
Parameters used for LoD calculations.

**Figure 1.**
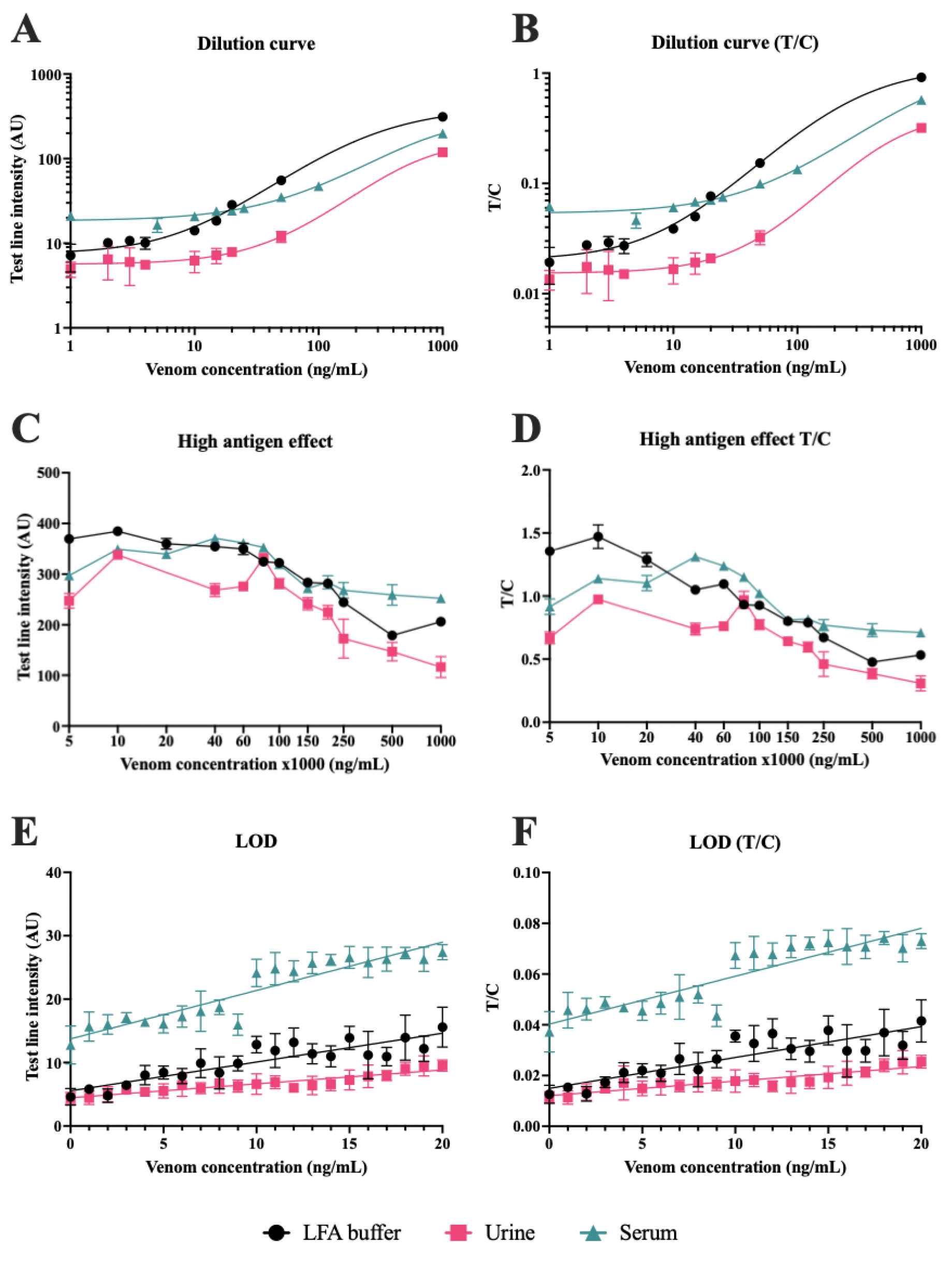
Dilution curves of sandwich pair 86-14 and 86-11 as measured with LFAs. A&B) B. atrox whole venom was dissolved in LFA running buffer at various concentrations and measured in duplicates in LFAs to assess the concentration-signal correlation of the test. Both the test line intensities as quantified with a commercial reader and the test line intensity to control line intensity (T/C) ratios are shown here. C&D) LFA running buffer was spiked with high concentrations of B. atrox whole venom and measured in triplicates in LFAs to assess the presence of hook effect. The concentrations displayed on the axis should be multiplied by 1,000 to get the actual concentration. E&F) Low concentrations of B. atrox venom were dissolved in LFA running buffer, pooled human serum, and pooled human urine, respectively, and measured in six replicates in LFAs to assess the LoD of the test.

### 3.3 Antigen identification

To identify the antigens recognised by the sandwich pair, the antibodies 86-14 and 86-11 were screened against *B. atrox* venom proteins using electrospray ionisation mass spectrometry (ESI-MS) to look for binding partners. These mass spectra were acquired under ‘soft’ ionisation conditions, which allow non-covalent interactions to be preserved and transferred into the gas phase. Prior to screening, *B. atrox* venom was fractionated by size exclusion chromatography (SEC) to separate the venom components by mass. This was done to generate venom protein mixtures, which were less complex than the whole venom, to make antigen identification easier, as well as to exchange the proteins into a spray solution that was appropriate for ESI-MS. **Figure 2A** shows the SEC profile of the venom, which contained multiple protein peaks (labelled in the chromatogram), which corresponded to proteins of different masses. The size exclusion fractions for these peaks were mixed with the two antibodies in a 1:1 (volume:volume) ratio to screen for binding. Both 86-11 and 86-14 antibodies only bound to venom proteins from SEC fraction four (elution volume 15-16 mL) of *B. atrox* venom. To better understand the nature of these antibody antigen interactions, 86-14 and 86-11 were titrated against *B. atrox* SEC fraction four. **Figure 2B & C** show the native mass spectra of both antibodies prior to mixing with the venom fraction, where the most abundant charge states were 24_^+^_ and 25^+^ for 86-11 and 86-14, respectively. **Figure 2D & E** shows the mass spectra of the antibodies mixed in a 1:1 (volume:volume) ratio with a diluted (one in five) *B. atrox* SEC fraction four. Within these spectra, two prominent charge state distributions were detected within the *m/z* region 5400 to 7600 for 86-11 and 86-14. The difference in masses between the charge state series corresponded to the antibodies complexed with a 23.3 kDa protein (168.1 kDa vs. 144.8 kDa for 86-11, and 169.1 kDa 146.5 kDa for 86-14). When mixed with the undiluted venom fraction, the most prominent charge state series have masses corresponding to the antibodies bound to 46.6 kDa of antigen (**Figure 2F & G**). Taken together, these titration experiments indicate that the antibodies are binding two 23.3 kDa venom proteins. To further test these observations, ions corresponding to antibody complexes with two toxins were isolated and fragmented by tandem MS (MS/MS) and subjected to collisional energy to dissociate the antigens.

**Figure 2.**
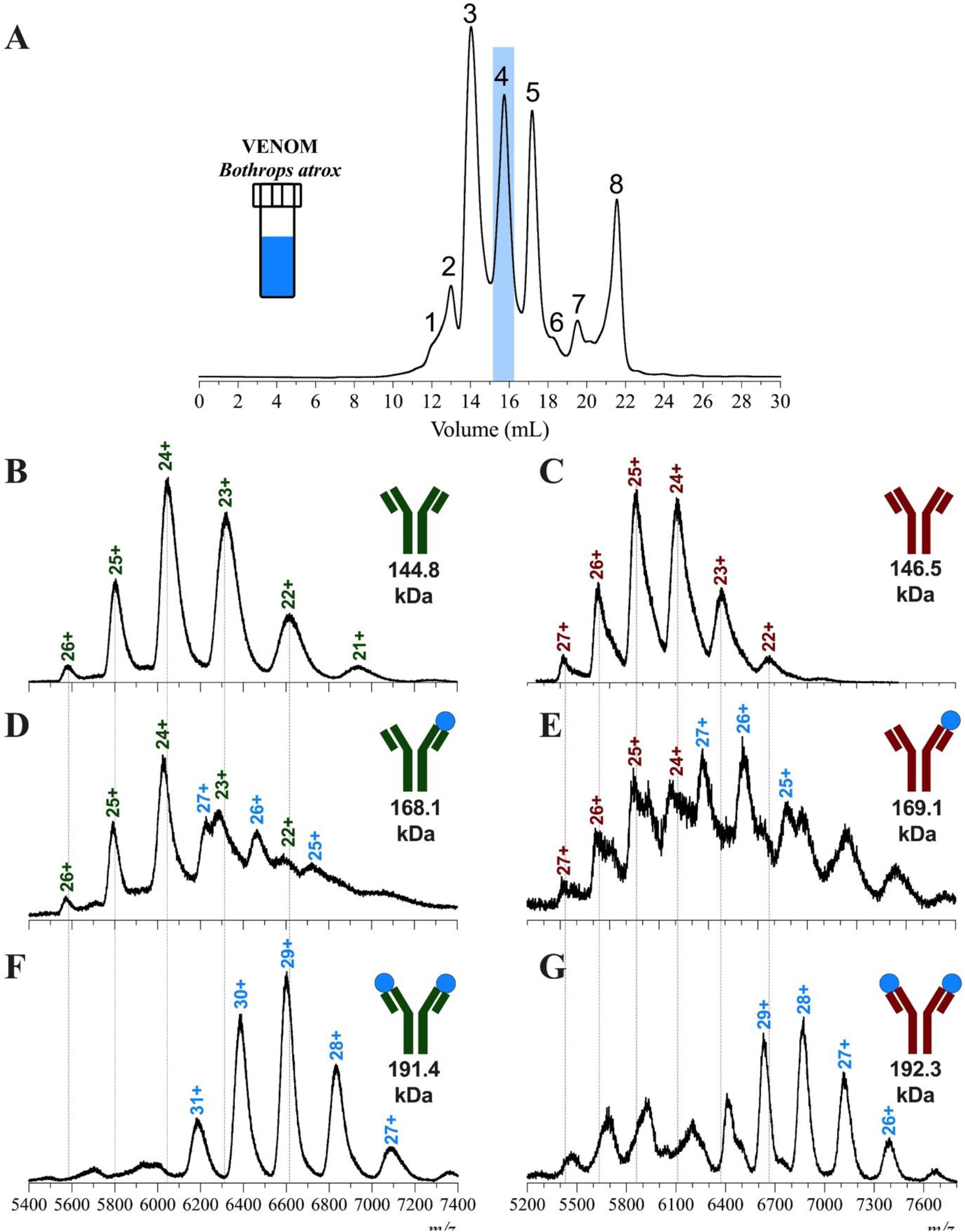
Size exclusion and Mass spectrometry for the titration of B. atrox venom against the monoclonal antibodies, 86-11 and 86-14. A) The size exclusion chromatogram of the B. atrox venom, with the peak corresponding to the antigen fraction highlighted in blue. Native mass spectra of B) antibody 86-11 (green) and antibody 86-14 (red), C) antibody 86-11 prior to mixing. The mass spectra for antibodies 86-14 and 86-11 mixed with the five times diluted (D & E, respectively) and undiluted (F & G, respectively) fraction from the SEC of B. atrox venom.

For the MS/MS experiments, the 28^+^ charge states of the antibody toxin complexes were isolated and subjected to collisional energy. This was done to dissociate the antigens from the antibody to accurately calculate their masses. **Figure 3** shows the MS/MS spectra of each antibody:toxin complex after the application of collisional energy. For both antibodies (**Figure 3A & B**), the ejected protein from the complexes had the mass of 23.3 kDa. A negative control was included to check the fragmentation patterns of only the 86-11 and 86-14 antibodies (**Error! Reference source not found**.). The data from these spectra show that the only antigen ejected has a mass of 23.3 kDa, and that this mass did not correspond to the fragmentation of the antibody. Taken together, the ESI-MS data presented in **Figure 2** and **Figure 3** show that the antibodies bind specifically to the 23.3 kDa toxin twice. The masses of the ejected toxins are within the expected sequence mass range of type I snake venom metalloproteases. However, further experiments are required to confidently identify the antigen.

**Figure 3.**
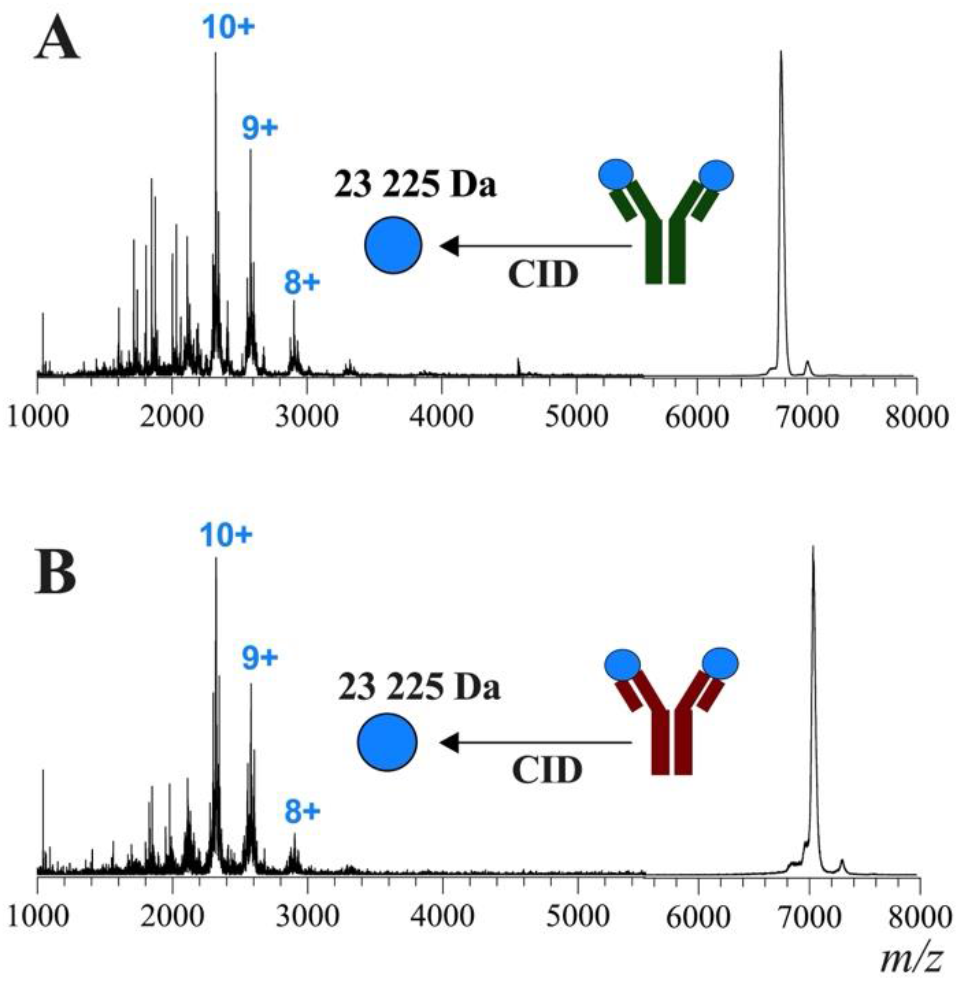
Tandem MS experiments for the antibody:(2)toxin complexes. Collision induced dissociation (CID) of antibody 86-11 (green) and antibody 86-14 (red) complexed with the antigens are shown in spectra A) and) B, respectively.

### 3.4 Assay characterisation

To further characterise the LFA, it was screened against the panel of 21 venoms used in the ELISAs, as well as an additional 24 venoms to determine which venoms were recognised (see **SI Table S1**). The results indicate that the assay is specific towards the venoms of *Bothrops* species from both Brazil and some of its neighbouring countries, while it cannot detect venoms from other viperids, such as *Crotalus* and *Lachesis* spp. (**Figure 4**).

**Figure 4.**
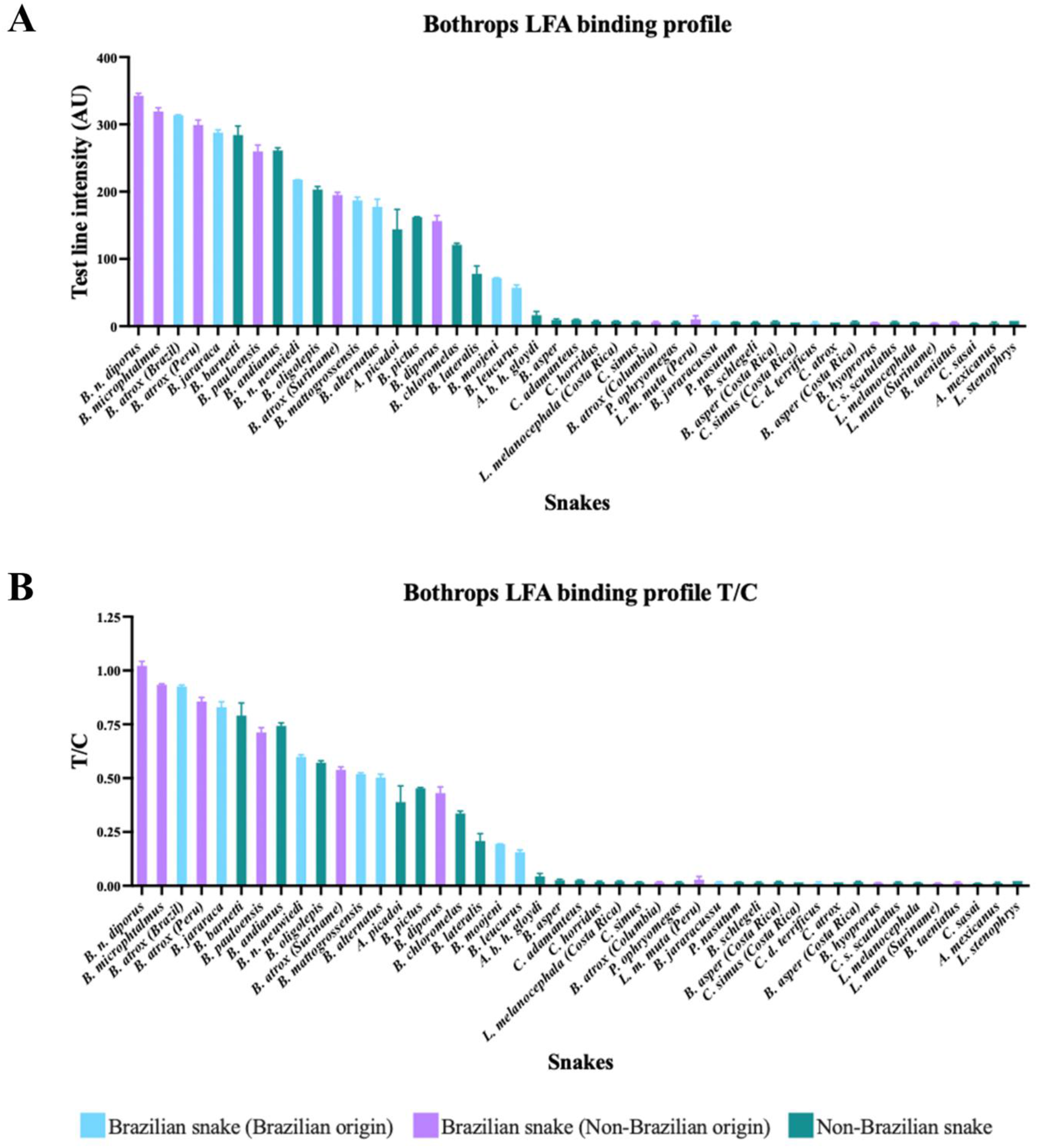
Venom recognition of the sandwich pair 86-14 and 86-11 in LFAs, when tested with 1000 ng/mL whole venom dissolved in LFA running buffer. LFA strip signals were quantified with a reader, and the test line signal to control line signal ratio (T/C ratio) is displayed here. Blue indicates that venom from Brazilian snake specimens were used. Purple indicates that the snake species is found in Brazil, but that the venom was extracted from a non-Brazilian specimen or that the origin of the specimen is unknown. Green indicates that the species is not found in Brazil.

Generally, LFAs are compatible with several different sample matrices. However, this requires careful optimisation to prevent matrix effects/false positives. To evaluate the effect of different sample matrices on venom detection by our antibodies, we spiked pooled human urine samples, pooled human serum samples, pooled human plasma (containing heparin) samples, and pooled human plasma (containing EDTA) samples with various amounts of *B. atrox* venom and detected the venom using both ELISAs and LFAs (**Figure 5** and **Error! Reference source not found**.). The results in urine were comparable to the results in running buffer, both in terms of the low occurrence of false positives on negative tests and high signals on positive tests. In LFAs, the serum samples generally elicited lower signal intensities than the other types of samples. Plasma samples with heparin had the highest signals on negative LFAs and medium signals on positive control LFAs.

**Figure 5.**
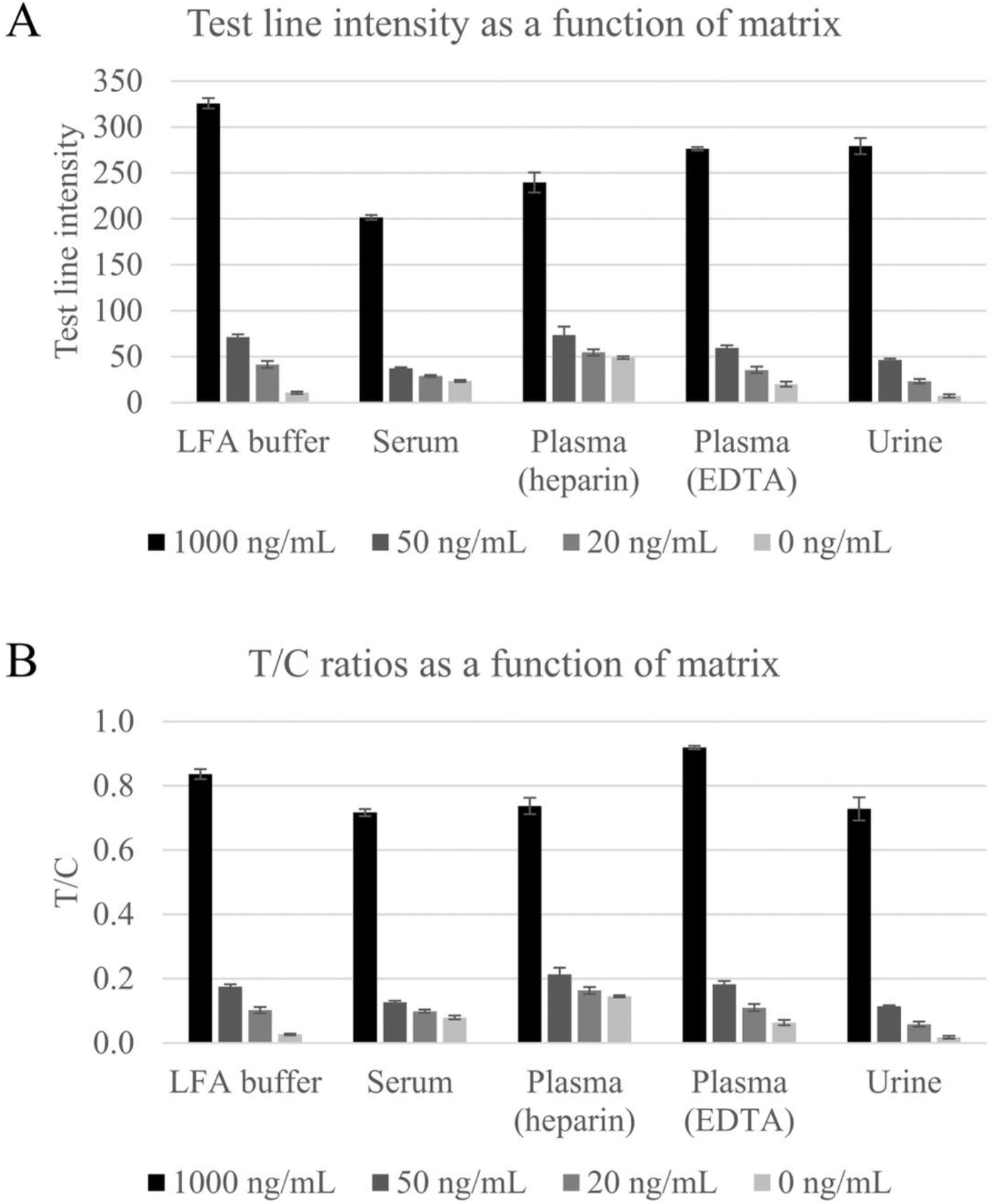
Comparison of matrix effects in LFAs. B. atrox venom was diluted in LFA running buffer, pooled human serum, pooled human plasma (containing heparin), pooled human plasma (containing EDTA), and pooled human urine. The antigen in the dilution series were detected with the LFA, and the test line intensities A) and T/C ratios B) are displayed here. The results shown are the averages of triplicates.

To assess whether compounds that might be found in human sample matrices could cause false positives in the LFA, serum and urine samples were spiked with a series of compounds (the individual concentrations of which can be seen in **SI S2**) and tested with LFAs in the absence of antigen. Additionally, the influence of sample pH on the presence of false positives was assessed. The results of the experiment can be seen in **Figure 6**. Urine samples generally yielded lower test line signals and higher control line signals than serum samples. The only tested compounds to cause false positives were glycine and heparin, and the only conditions to cause false positives were pHs 4.0, 5.0, and 5.5, and in all cases these false positives only occurred when tested in serum, indicating that serum causes matrix effects for the reported LFA.

**Figure 6.**
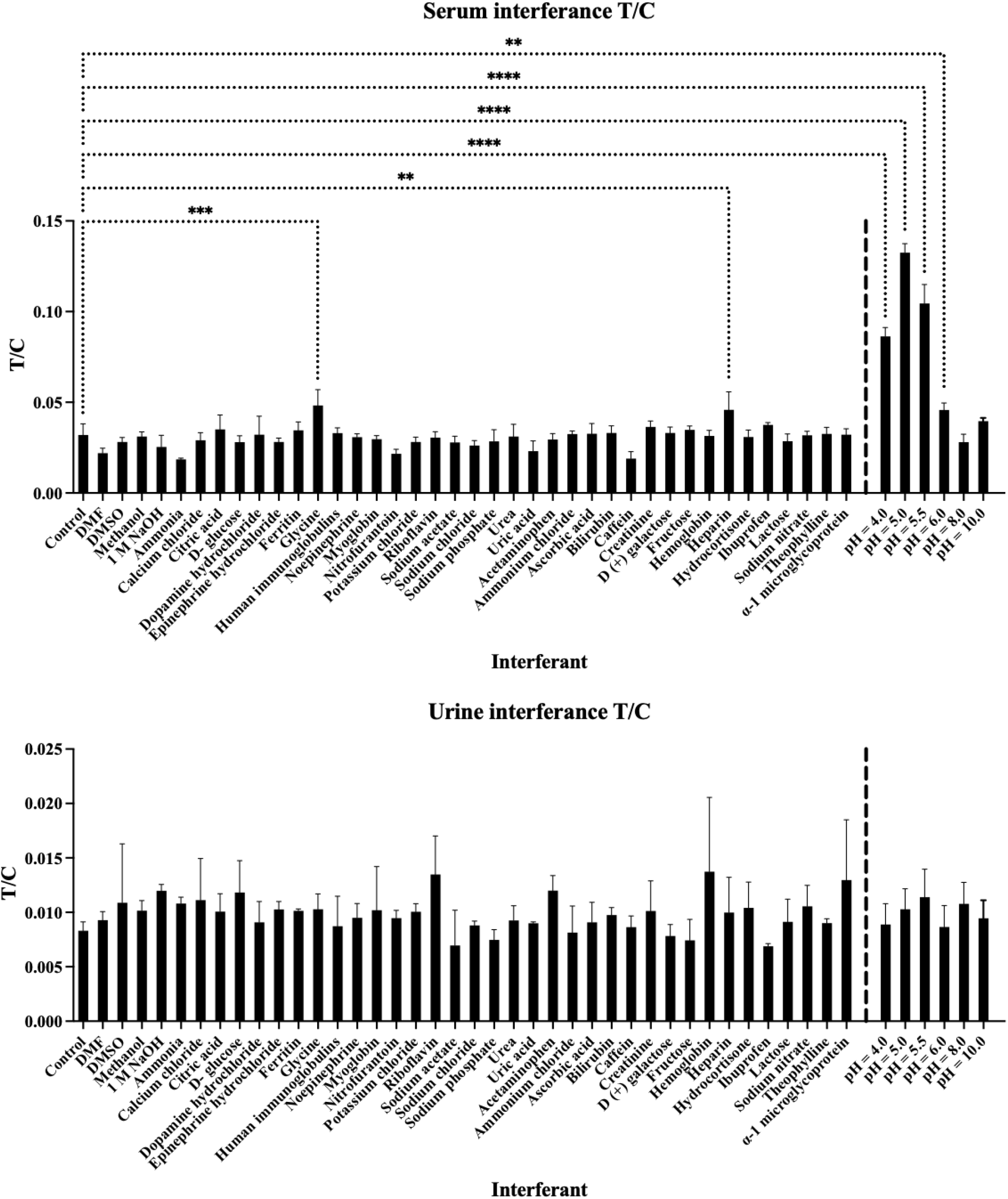
Screening of potential interferants in LFAs. Serum and urine samples were spiked with different compounds and tested in LFAs in the absence of antigen to investigate whether the compounds could cause false positives. In addition, the solvents (water, DMF, DMSO, and methanol) used for dissolving the compounds were tested. Serum and urine at their natural pHs, and pH-adjusted serum and urine samples were also tested.

### 3.5 Assay stability

It was observed that the conjugated antibodies were not stable at room temperature for extended periods of time (data not shown). This could decrease the utility of the assay intended for use in the tropics. To mitigate this thermal instability of the conjugated antibodies, it was decided to lyophilise them, and store them at 4 °C until use, at which point they were reconstituted in the running buffer and the sample to be tested. The antibodies were lyophilised in six different solutions, consisting of either bovine serum albumin (BSA) or casein as blocking agents, and with either trehalose, sucrose, or mannitol as stabilising agents. The lyophilised antibodies were then used to run LFAs in which a dilution curve of venom was tested. It was determined that the solutions blocked with casein and stabilised with either trehalose or sucrose, respectively, yielded the best results, as no false positives were observed in negative samples, while a comparatively strong test line signal was retained in positive samples (data not shown). Conjugated antibodies lyophilised in the casein-trehalose and casein-sucrose solutions were tested further: The lyophilised antibodies were stored at -20 °C, 4 °C, room temperature (roughly 22 °C), and 37 °C; and tested either immediately after lyophilisation, after one week, two weeks, four weeks, and eight weeks, on venom concentrations of 500 ng/mL and 0 ng/mL. For the 500 ng/mL LFAs, a decrease in test line intensity occurred within the first two weeks, after which the signal stabilised (**Figure 7**). The LFAs run with antibodies lyophilised in the sucrose solution had slightly higher test line intensities at week eight than the LFAs run with antibodies lyophilised in the trehalose solution (**Figure 7**). Taken together, these results indicate that it is possible to extend the shelf-life of the conjugated antibodies by lyophilising them, thereby potentially extending the shelf-life and overall utility of the LFA.

**Figure 7.**
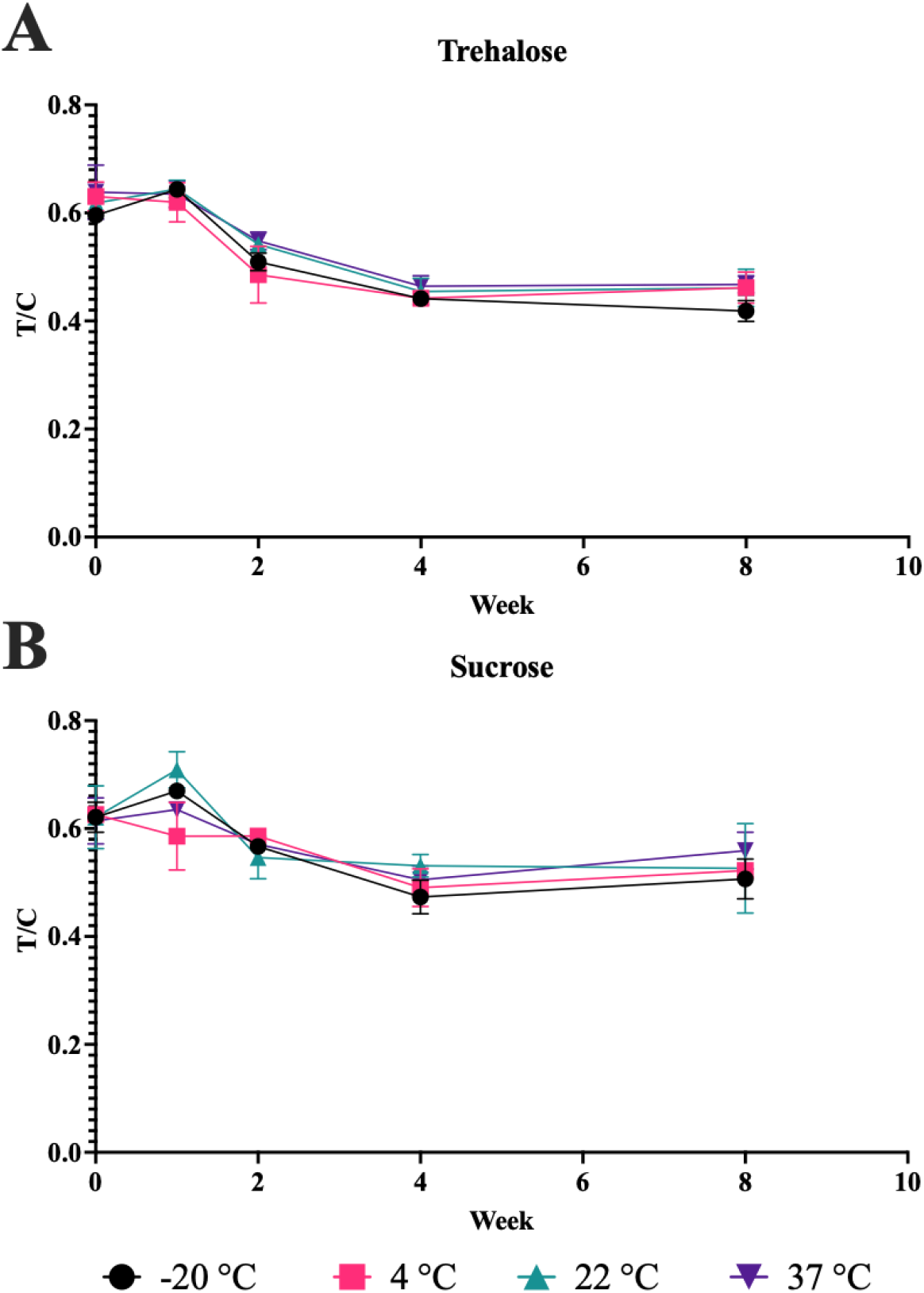
T/C ratios of LFAs utilising lyophilised antibodies stored under different conditions to detect 500 ng/mL whole venom from B. atrox. The gold-conjugated and biotinylated antibodies were lyophilised in solutions containing either A) trehalose or B) sucrose. The lyophilised reagents were then stored at either -20 °C, 4 °C, 22 °C, or 37 °C for between 0 and 8 weeks, before being used in LFAs to detect 500 ng/mL B. atrox venom dissolved in LFA running buffer.

## 4 Discussion

The LFA prototype presented here is the first report of the use of monoclonal antibodies in an LFA to detect *Bothrops* venom. The LoD of the prototype, when read with a reader, was 8.0 ng/mL in serum samples and 9.5 ng/mL in urine samples, with a visual LoD of approximately 25 ng/mL in buffer. This LoD would possibly be sufficient to detect venom in most patient samples, as a literature study found that reported venom concentrations in patient samples across species varied from < 1 ng/mL to > 1000 ng/mL, and that venom concentrations were generally higher after bites caused by vipers than after bites caused by elapids [27]. Two case studies of patients with bothropic envenomings revealed serum venom concentrations of 33.7 ng/mL and 62.6 ng/mL five hours post-bite, respectively, well above the LoDs reported here [23,24]. However, it is unknown whether these venom concentrations are representative of average bothropic envenomings. Generally, venom concentrations in patient samples are expected to vary depending on the sample matrix (e.g., urine or serum), time since the bite occurred, the body size of the patient, and other parameters. If the LoD of the prototype proves to be too high to detect venom in a certain sample type or at a certain time (e.g., in a urine sample immediately after the bite), it might be possible to improve the LoD by enriching the venom from the sample, or alternatively to measure another sample type (e.g., a wound swab or serum sample) [28,29]. Conversely, if the antigen concentration in the sample is high enough to bring the assay into the hook effect range, it might be possible to dilute the sample.

One limitation of this study is that the LFA was not evaluated on real patient samples. Therefore, it remains to be seen to what extent the LFA is capable of detecting venom in real samples and whether any untested compounds interfere and cause false positives or false negatives. Two compounds that could potentially cause issues are antivenom and biotin. Antivenom is often added to snakebite patient blood samples to prevent clotting, and it is likely that the polyvalent antivenom antibodies can block the epitopes recognised by the monoclonal antibodies employed in the LFA. This potential issue might be alleviated by measuring serum or plasma samples derived from blood samples collected before antivenom administration or by using different sample types. As for biotin, it has recently been suggested that overconsumption of biotin by the general population might lead to elevated biotin levels in plasma, which could potentially interfere with immunoassays relying on biotin-binding, such as the gRAD [30].

A study showed that 4.24% of snakebite victims in Brazil received two or more kinds of antivenom, and that 10.5% of patients bitten by *Bothrops* spp. received polyvalent antivenom [31]. Administration of polyvalent or multiple antivenoms might indicate a lack of confidence in the identification of the type of snakes involved in snakebite accidents. This notion could be supported by another study of 1063 snakebites in Brazil, in which it was found that only 44% of snakebites were identified at the genus level [32]. Lack of confidence in the identification of the snake involved in an accident is potentially problematic, as it has been demonstrated that a delay in treatment, either due to insufficient or incorrect administration of antivenom, for *Crotalus* bites in Brazil led to an increased risk of acute renal failure for the patients [13]. Additionally, it has been argued that there is a systematic lack of training of healthcare professionals in clinical snakebite management in certain states in Brazil [33,34]. Thus, snakebite diagnostic tools could potentially help mitigate the risk of inappropriate antivenom administration and promote the use of monovalent antivenoms, especially in cases where the treating personnel is not trained in clinical snakebite management. In the future, the LFA might also be used to confirm envenoming prior to administration of upcoming first-line-of-defence drugs, if drug candidates such as varespladib and marimastat are approved for treatment of snakebite envenoming [35]. This could in turn improve patient outcome even further.

Taken together, the data presented here constitute a proof-of-concept for a rapid diagnostic test for *Bothrops* envenomings. Potentially, a diagnostic tool, such as the LFA presented here, could be used to confirm or disprove suspected *Bothrops* envenomings, thereby guiding the choice of antivenom not only for the 10.5% of patients with bothropic bites who receive polyvalent antivenom but also for the 34.4% of patients with crotalic bites who receive either bothropic or bothro-crotalic antivenom [31].

## 5 Conclusions

Here, we have presented a prototype LFA capable of distinguishing the venoms of several *Bothrops spp*. from the venoms of non-*Bothrops spp*. The LFA had an LoD of 8 ng/mL in serum and 9.5 ng/mL in urine, when read with a commercial reader, and a visual LoD below 25 ng/mL. In future studies, we plan to further improve the LFA’s sensitivity, prior to evaluating its performance on patient samples. It is possible that the LFA could empower healthcare providers who have limited or no experience in clinical snakebite management to diagnose snakebite victims more confidently. This might be especially valuable in remote clinics and as a support to ongoing efforts to decentralise antivenom in the Brazilian Amazon, thereby bringing treatment closer to those who need it most and ameliorating the burden of the highly neglected disease of snakebite envenoming.

## 6 Acknowledgments

The authors would like to thank Prof. Bruno Lomonte of Instituto Clodomiro Picado at Universidad de Costa Rica, Dr.s Alfonso Zavaleta and Maria Salas of Cayetano Heredia University, and Dr. Soledad Bustillo of Universidad Nacional del Nordeste for their kind donations of lyophilised venoms. The authors would also like to thank Prof. Ana Moura da Silva of Instituto Butantan, Profs. Wuelton Monteiro, Marco Sartim, and Lisele Brasileiro of Amazonas State University, and Prof. Manuela Pucca of the Federal University of Roraima for good scientific discussions.

## Author contributions

CK, AHL, JAJ, SHD, AMH, and RUWF conceived the study. CK, SHD, AMH, RUWF, PDK, SBB, JAH, and IO carried out the experiments and analysed the data. CK, JAJ, IO, and JAH drafted the manuscript. All authors revised and reviewed the manuscript.

