## Supplemental information for "Prototyping of a lateral flow assay based on monoclonal antibodies for detection of *Bothrops* venoms"


| | | |
|---|---|---|
| **Table S1** | List of venoms used in the study | S2 |
| **Table S2** | List of compounds, solvents, & concentrations for interference study | S5 |
| **Figure S1** | Antibody binding profiles of 21 venoms in ELISA | S7 |
| **Figure S2** | Sandwich pair binding profiles of 21 venoms in ELISA | S8 |
| **Figure S3** | Photos of lateral flow assays testing a dilution curve of venom | S9 |
| **Figure S4** | Tandem MS experiments showing antibody fragmentation | S9 |
| **Figure S5** | Matrix effects in ELISA | S10 |

***Table S1.*** *List of venoms used in this study, sorted by genus, species, origin, and manufacturer.*

| Genus | Species and subspecies | Origin | Manufacturer/donor |
|---|---|---|---|
| *Agkistrodon* | *bilineatus howardgloydi* | Costa Rica (Guanacaste) | Prof. Bruno Lomonte |
| *Atropoides* | *Mexicanus** | Honduras | Prof. Bruno Lomonte |
| *Atropoides* | *picadoi* | Costa Rica | Prof. Bruno Lomonte |
| *Bothriechis* | *lateralis* | Costa Rica | Prof. Bruno Lomonte |
| *Bothriechis* | *schlegelii* | Costa Rica (Atlantic + pacific) | Prof. Bruno Lomonte |
| *Bothrops* | *alternatus* | | Kentucky Reptile Zoo, cat# BAL |
| *Bothrops* | *andianus* | Peru | Dr.s Alfonso Zavaleta & Maria Salas |
| *Bothrops* | *asper* | Costa Rica (Carribean) | Prof. Bruno Lomonte |
| *Bothrops* | *asper* | Costa Rica (pacific) | Prof. Bruno Lomonte |
| *Bothrops* | *asper* | | Latoxan |
| *Bothrops* | *atrox* | Brazil | Latoxan, cat#L1210A |
| *Bothrops* | *atrox* | Brazil, Peru, French Guyana | Latoxan, cat#L1210 |
| *Bothrops* | *atrox* | Columbia | Kentucky Reptile Zoo, cat# BAt-C |

| | | | |
|---|---|---|---|
| *Bothrops* | *atrox* | Peru | Dr.s Alfonso Zavaleta & Maria Salas |
| *Bothrops* | *atrox* | Surinam | Kentucky Reptile Zoo, cat# BAt-S |
| *Bothrops* | *barnetti* | Peru | Dr.s Alfonso Zavaleta & Maria Salas |
| *Bothrops* | *chloromelas* | Peru | Dr.s Alfonso Zavaleta & Maria Salas |
| *Bothrops* | *diporus* | Argentina | Dr. Soledad Bustillo |
| *Bothrops* | *diporus*** | | Latoxan, cat# L1267 |
| *Bothrops* | *hyoprorus**** | Peru | Dr.s Alfonso Zavaleta & Maria Salas |
| *Bothrops* | *jararaca* | | Latoxan, cat# L1211 |
| *Bothrops* | *jararacussu* | | Latoxan, cat# L1256 |
| *Bothrops* | *leucurus* | | Latoxan, cat# L1255 |
| *Bothrops* | *mattogrossensis* | | Latoxan, cat# L1248 |
| *Bothrops* | *microphthalmus* | Peru | Dr.s Alfonso Zavaleta & Maria Salas |
| *Bothrops* | *moojeni* | | Latoxan, cat# L1254 |
| *Bothrops* | *neuwiedi neuwiedi* | | Latoxan, L1213 |
| *Bothrops* | *oligolepis* | Peru | Dr.s Alfonso Zavaleta & Maria Salas |
| *Bothrops* | *pauloensis* | | Latoxan, cat# L1265 |

| | | | |
|---|---|---|---|
| *Bothrops* | *pictus* | Peru | Dr.s Alfonso Zavaleta & Maria Salas |
| *Bothrops* | *taeniatus* | Peru | Dr.s Alfonso Zavaleta & Maria Salas |
| *Cerrophidion* | *sasai* | Costa Rica | Prof. Bruno Lomonte |
| *Crotalus* | *adamanteus* | | Kentucky Reptile Zoo, cat# EDB |
| *Crotalus* | *atrox* | | Kentucky Reptile Zoo, cat# CAt |
| *Crotalus* | *durissus terrificus* | Brazil | Latoxan, cat# L1261B |
| *Crotalus* | *horridus* | | Kentucky Reptile Zoo, cat# CH |
| *Crotalus* | *scutulatus scutulatus* | | Kentucky Reptile Zoo, cat# CSS-T |
| *Crotalus* | *simus* | Costa Rica | Prof. Bruno Lomonte |
| *Crotalus* | *simus* | | Latoxan |
| *Lachesis* | *melanocephala* | Costa Rica | Prof. Bruno Lomonte |
| *Lachesis* | *melanocephala* | Costa Rica (pacific) | Prof. Bruno Lomonte |
| *Lachesis* | *muta* | Suriname | Latoxan, cat# L1290B |
| *Lachesis* | *muta muta* | Peru | Dr.s Alfonso Zavaleta & Maria Salas |
| *Lachesis* | *stenophrys* | Costa Rica (Caribbean) | Prof. Bruno Lomonte |

| | | | |
|---|---|---|---|
| *Porthidium* | *nasutum* | Costa Rica | Prof. Bruno Lomonte |
| *Porthidium* | *ophryomegas* | Costa Rica | Prof. Bruno Lomonte |

*Also known as *Metlapilcoatlus mexicanus* and *Bothrops nummifer mexicanus*.

**Purchased as *Bothrops neuwiedi diporus*.

***Also known as *Bothrocophias hyorprora*.

***Table S2***. *List of compounds tested for interference, the compounds' solvents, and the concentrations they were tested at.*

| Interferant | Solvent | Purity | Concentration mg/mL | Manufacturer | Product number | Batch number |
|---|---|---|---|---|---|---|
| **1 M NaOH** | - | - | - | VWR Chemicals | 31.624.290 | 17L134002 |
| **a-1 Microglycoprotein** | Sample | - | 0.04 | Lee Biosolutions | 126-11 | 04B5042 |
| **Acetaminophen** | Water | 100 | 0.40 | SIGMA | A7085-100g | SLBM5923V |
| **Ammonia** | Methanol | 100 | 2.00 | | 779423 | |
| **Ammonium Chloride** | Water | - | 10.00 | Sigma | A9434-500g | BCCB8234 |
| **Ascorbic Acid** | Water | - | 0.30 | Sigma Aldrich | PHR1008 | LRAC2886 |
| **Bilirubin** | Water | - | 0.003 | Sigma | B4126-1g | SLBN3618V |
| **Caffein** | Water | - | 1.04 | Sigma Aldrich | C0750 | BCBQ5381V |
| **Calcium Chloride** | Water | 100 | 1.00 | MERCK | 1023821000 | A0263182 144 |
| **Citric Acid** | Water | 100 | 1.00 | SIGMA | 1.00244.1000 1kg | BCBZ1757 |
| **Creatinine** | Water | - | 2.00 | Sigma Aldrich | C4255-10mg | SLCF3194 |
| **D (+) Galactose** | Water | 100 | 0.80 | SIGMA | G0750 5g | BCBX1697 |
| **D-Glucose** | Water | 100 | 15.00 | SIGMA | G8270 10g | SZBF0610V |
| **DMF** | - | - | - | MERCK | 1.029.370.500 | 1012737 |
| **DMSO** | - | - | - | SIGMA | D8414 | SHBJ5443 |

| | | | | | | |
|---|---|---|---|---|---|---|
| **Dopamine Hydrochloride** | Water | - | 0.066 | Sigma | H8502-5g | BCBQ4375V |
| **Epinephrine Hydrochloride** | Water | - | 0.066 | Sigma | E4642-5g | MKBS9064V |
| **Ferritin** | Water | 95 | 0.066 | - | 270-40 1mg | - |
| **Fructose** | Water | 100 | 1.00 | SIGMA | PHR1002 1g | LRAB2150 |
| **Glycine** | Water | 100 | 5.00 | MERCK MILLIPORE | | VP708301 543 |
| **Hemoglobin** | Water | - | 10.00 | Sigma | H7379-1g | SLBR9638V |
| **Heparin** | Water | - | 0.31 | Sigma | H0878-100KU | SLBN9256V |
| **Human Immunoglobulins** | Water | 100 | 0.25 | Produced in-house | N/A | - |
| **Hydrocortisone** | Water | - | 0.0906 | Sigma | H0888-1g | SLBL4101V |
| **Ibuprofen** | Water | - | 5.00 | Sigma-Aldrich | PHR1004 | LRAC5691 |
| **Lactose** | Water | 100 | 0.10 | FLUKA (SIGMA) | 17814 1kg | 1167331 |
| **Methanol** | - | - | - | - | M1775-1GA | - |
| **Myoglobin** | Water | 100 | 0.066 | - | 431-11 1mg | - |
| **Nitrofurantoin** | DMF | 100 | 0.60 | SIGMA | N7878 10g | MKBW0733V |
| **Norepinephrine** | Water | - | 0.066 | Sigma | A9512-250 mg | SLBJ1569 |
| **pH = 4.0** | - | - | - | - | - | - |
| **pH = 5.0** | - | - | - | - | - | - |
| **pH = 5.5** | - | - | - | - | - | - |
| **pH = 6.0** | - | - | - | - | - | - |
| **pH = 8.0** | - | - | - | - | - | - |
| **pH = 10.0** | - | - | - | - | - | - |
| **Potassium Chloride** | Water | 100 | 15.00 | SIGMA | PHR1329 | LRAB7667 |
| **Riboflavin** | Water | 100 | 0.01 | SIGMA | R4500 5g | WXBC7130V |
| **Sodium Acetate** | Water | 100 | 0.25 | SIGMA | S2889 250g | SLBX8569 |
| **Sodium Chloride** | Dissolved in sample | 100 | 40.00 | FISHER CHEMICALS | S/3160/65 | 1873920 |
| **Sodium Nitrate** | Water | 100 | 0.10 | SIGMA | S5506 250g | MKCH0828 |

| | | | | | | |
|---|---|---|---|---|---|---|
| **Sodium Phosphate** | Water | 96 | 5.00 | SIGMA | 342483 25g | MKCB7570 |
| **Theophylline** | Water | 100 | 0.10 | SIGMA | T1633 50g | MKBS5105V |
| **Urea** | Water | 100 | 40.00 | FLUKA (SIGMA) | 51465 1kg | 451840/1 |
| **Uric acid** | 1 M NaOH | 100 | 1.50 | SIGMA | U2625 | BCBP7874V |
| **Water (15 uL)** | - | - | - | - | - | - |

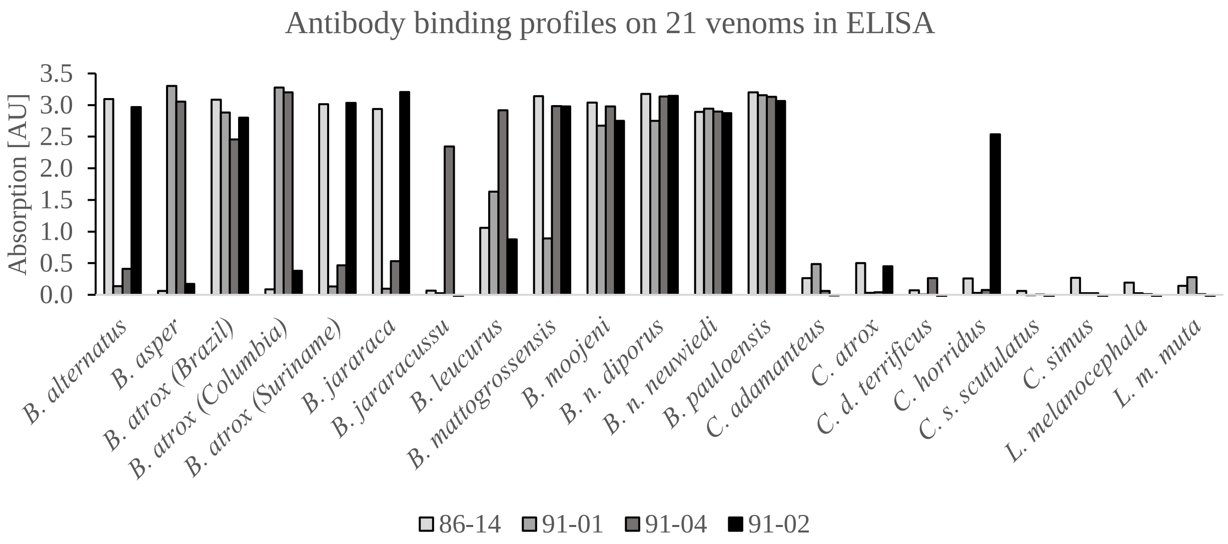


***Figure S1**. Binding profiles of antibodies 86-14, 91-01, 91-02, and 91-04 when screened against 21 venoms in an ELISA. Antibodies 86-14, 91-01, and 91-04 elicit strong signals when tested against several of the Bothrops venoms, but not when tested against Crotalus or Lachesis venoms. Antibody 91-02 also elicits strong signals on several Bothrops venoms, but unlike the other antibodies, it also elicits a strong signal on C. horridus venom.*

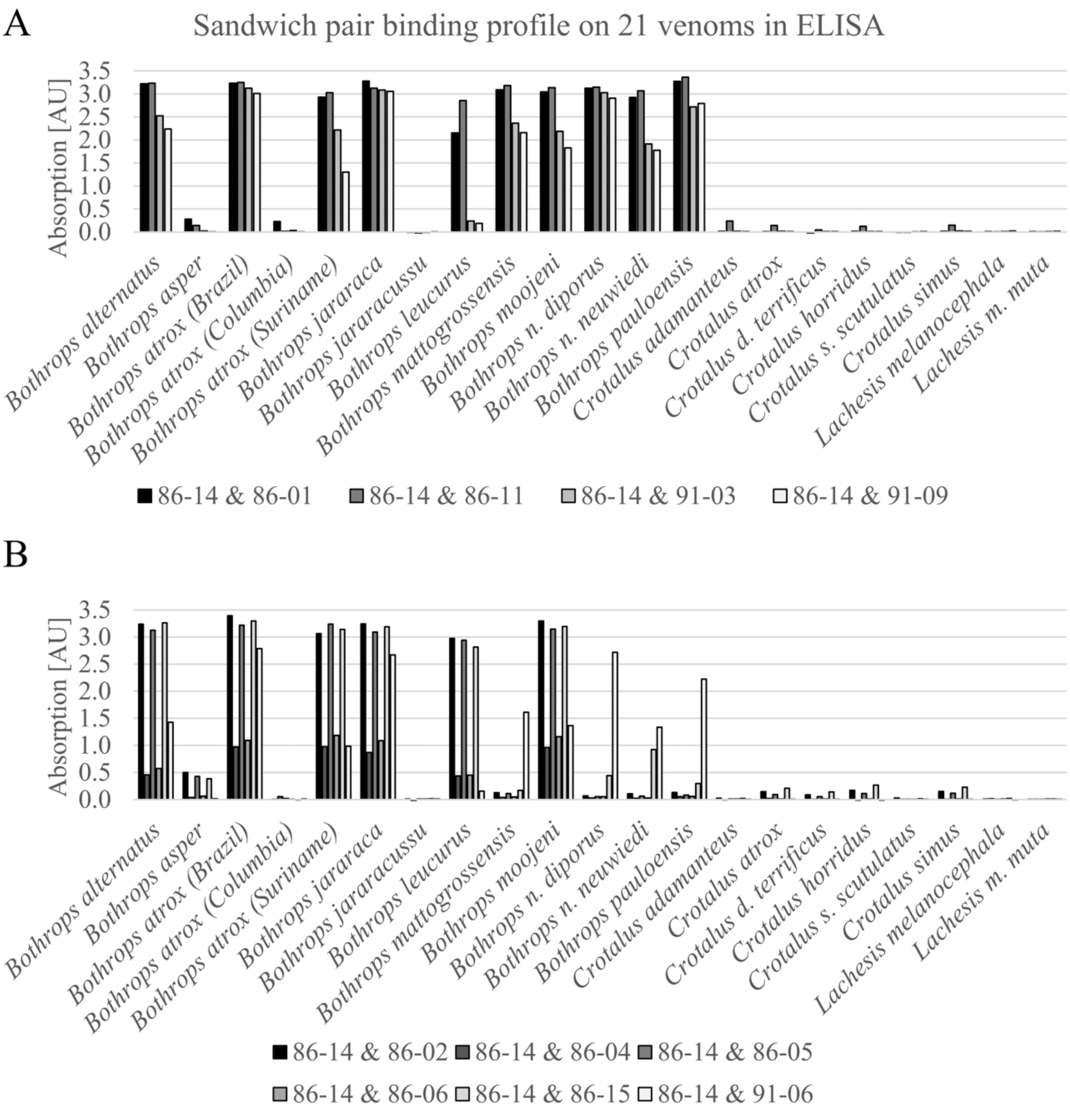


***Figure S2***. *Binding profiles of the ten sandwich pairs when screened in ELISA against 21 venoms. A) The sandwich pairs that were selected for further analysis based on their binding profiles. B) The sandwich pairs that were not selected for further analysis.*

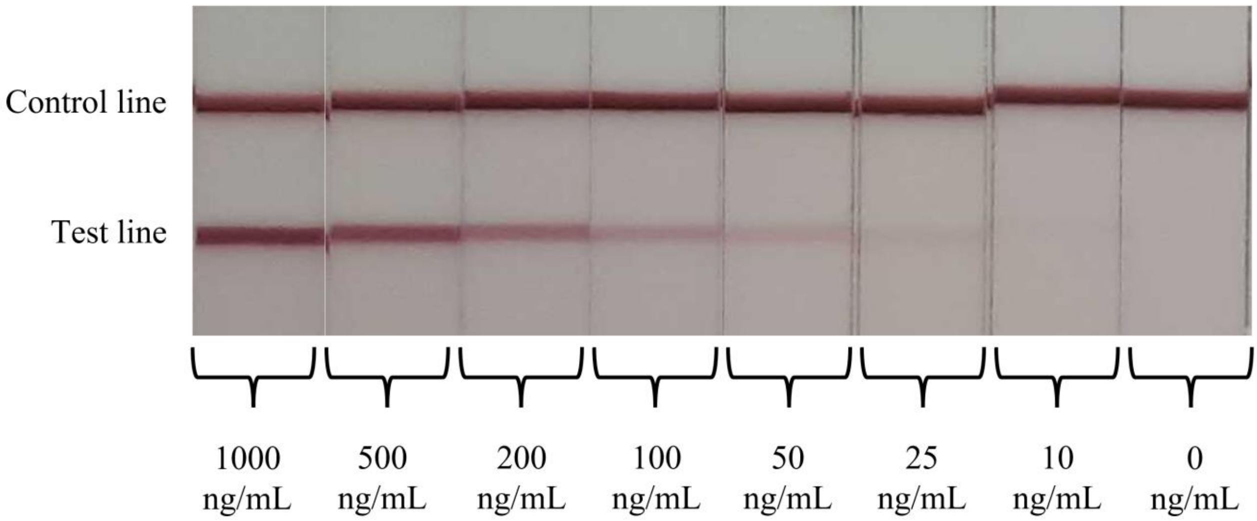


***Figure S3***. *Photos of lateral flow assays which were used to test a dilution curve of B. atrox venom dissolved in running buffer. An LoD of 10.3 ng/mL was achieved with a commercially available reader, while a visual LoD of 25 ng/mL could be detected by the naked eye.*

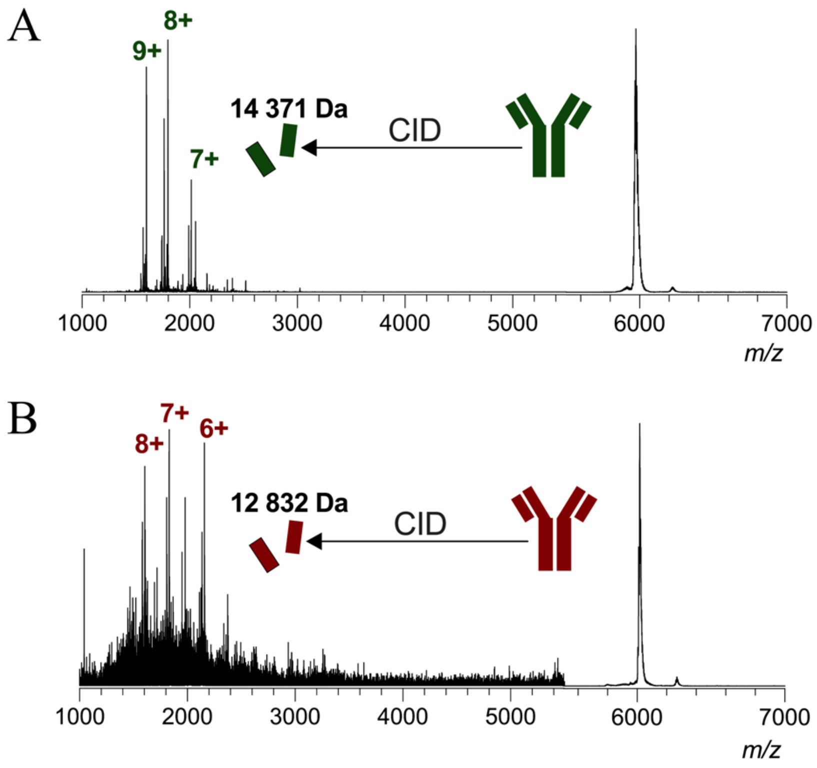


***Figure S4***. *Resulting spectra from tandem MS experiments for A) antibody 86-11 and B) 86-14, showing the fragmentation patterns of these two antibodies.*

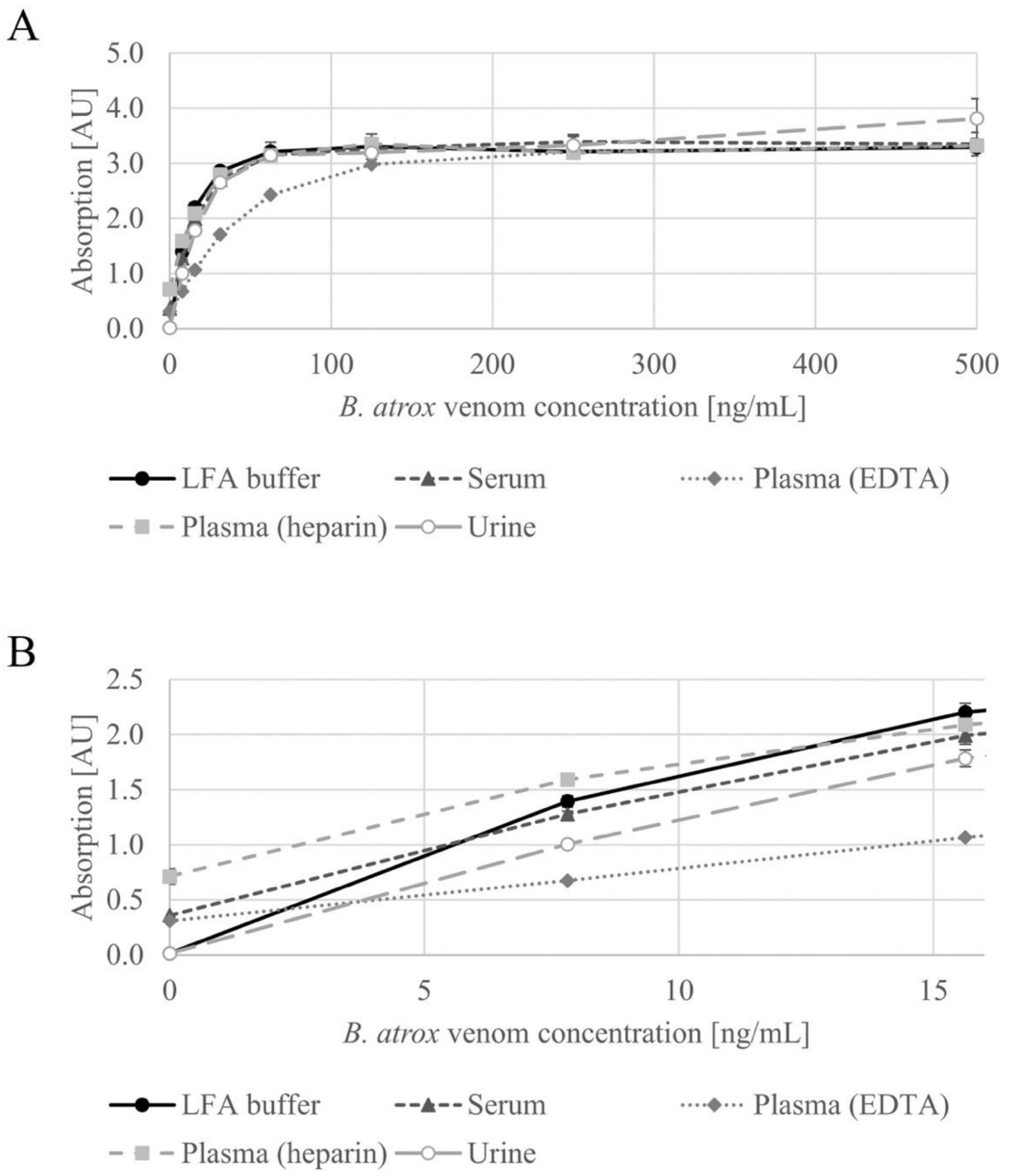


***Figure S5**. Matrix effect in ELISA. B. atrox venom was diluted in LFA running buffer, pooled human serum, pooled human plasma (heparin), pooled human plasma (EDTA), and pooled human urine. The dilution series were measured with ELISA, and the test absorbances at the different concentrations are shown in A). B) shows a subsection from A) of the signals corresponding to venom concentrations ranging from 0-16 ng/mL. This subsection reveals the different y-axis intercepts of the dilution curves, which in turn indicate different levels of false positive signal. It also shows differences in signal strength across the different matrices, in spite of the tested venom concentrations being the same. The results shown are the averages of duplicates.*
